# Hydrogel Microneedle Array-Based Transdermal Dressing System for Multiplexed Assessment and Combined Therapy of Chronic Wounds

**DOI:** 10.1101/2023.12.08.570882

**Authors:** Md Sharifuzzaman, Gauri Kishore Hasabnis, Sheikh Ahmed Abu Saleh, Leonard Siebert, Gregor Maschkowitz, Zeynep Altintas

**Affiliations:** Chair for Bioinspired Materials and Biosensor Technologies, Institute of Materials Science, Faculty of Engineering Christian-Albrechts-Universität zu Kiel / Kiel University, Kaiserstr. 2, 24143 Kiel, Germany; Chair for Functional Nanomaterials, Institute of Materials Science, Faculty of Engineering Christian-Albrechts-Universität zu Kiel / Kiel University, Kaiserstr. 2, 24143 Kiel, Germany; Institute for Infection Medicine, Christian-Albrechts-Universität zu Kiel / Kiel University, Brunswiker Str. 4, 24105 Kiel, Germany

**Keywords:** Conductive hydrogel-forming microneedles (HFMNs) array, Laser-scribed phase separation, Chronic wounds, transdermal dressing, Diagnostics and therapy, Correlation between wound exudate and interstitial fluid

## Abstract

Although recent wearable chronic wound (CWO) bandage technologies have opened up exciting opportunities for personalized CWO management, they still face significant obstacles due to the reliance on the wound bed exudate for sensing and delivering therapeutics. Flat, shallow, and desiccated wounds are difficult to collect wound exudate for sensing, and some wounds continuously exude, potentially washing delivered therapeutics out of the wound bed. Herein, we developed a hydrogel-forming microneedles (HFMNs) array-based multimodal transdermal dressing system that continuously monitors the on-site physiological conditions of CWOs in interstitial fluid (ISF) and offers healing capabilities. The unique polar array design enables the integration of six replaceable HFMNs sensing electrodes to target the desired wound-specific analytes in transdermal interstitial fluid (glucose, uric acid, pH, Na^+^, Cl^-^, K^+^, and temperature) based on their significance in reflecting the status of the CWOs. The hydrogel is composed of a biocompatible and swellable polymer - polyvinyl alcohol, and chitosan as a crosslinking agent, while the incorporation of MXene (Ti_3_C_2_T_x_) nanosheets as conductive nanofillers facilitates the formation of 3D polymer hydrogel networks via hydrogen bonding. Further coating and functionalization of poly(3,4-ethylenedioxythiophene): polystyrene sulfate (PEDOT: PSS) and graphene oxide through a laser-scribed phase separation (LSPS) process improves the electrical conductivity and in-vivo water stability of the HFMNs as a result of the larger and interconnected PEDOT-rich domains. Importantly, anti-inflammatory and antibacterial properties of the hydrogel prevent wound infection and promote skin wound healing. Through the potential correlation between wound-affected ISF and wound bed exudate, this method bridges conventional and implantable dressing systems for commercialization.

## 1. Introduction

Over 2 million people in Europe and 6.5 million people in the United States suffer from chronic nonhealing wounds, such as diabetic ulcers, nonhealing surgical wounds, burns, and venous-related ulcerations^[1]^, costing the health care system over $25 billion annually. Hemostasis, inflammation, proliferation, and remodeling are the four interdependent and contiguous phases of CWOs healing^[2]^. In general, wound exudate (WX) production is at its maximum during the inflammatory phase and declines as the lesion heals^[3]^. At each stage, the chemical composition (e.g., pH, glucose, uric acid, and ions) and physical parameters (e.g., temperature) of the wound environment change significantly, indicating the stage of wound healing and even the presence of infection^[4–8]^.

Planimetry is presently utilized in the clinical evaluation of wounds to qualitatively score characteristics such as slough reduction, granulation tissue formation, and reepithelialization^[9]^. Currently, quantitative profiling of biochemical parameters is typically limited to laboratory testing, such as enzyme-linked immunosorbent assays (ELISAs)^[10]^. Furthermore, several of the existing treatments, such as skin replacements, tissue transplants, mechanical wound psychotherapy, and others, can be helpful but often need surgery ^[1]^. Bacterial infection at the lesion site may result in tissue necrosis, by greatly impeding the healing process. A growing number of patients are being prescribed therapeutics, including both topical and systemic treatments, for chronic nonhealing lesions. An examination of the wound environment, nevertheless, reveals that an eschar generally causes a division of viable cells beneath the epidermis from its outermost layer.^[2]^ This necessitates the diffusion of therapeutics via the eschar in order to reach viable cells following topical application. Consequently, the topical delivery of medications results in a reduced local bioavailability compared to initial expectations.^[11]^

Despite the promising prospects that wearable smart dressings have presented^[12–15]^, the selection of the optimal site of action remains a prevalent issue for sensing systems and drug delivery objects utilized in wound care. The majority of present wearable dressing systems rely on stratum corneum-level WX with relatively limited and insufficient information. Flat, shallow, and desiccated wounds are difficult to collect WX for sensing, and some wounds continuously exude, potentially washing delivered therapeutics out of the wound site. On the other hand, the changes in the physiological surroundings of healthy cells beneath eschar may not be precisely reflected in the values obtained with topical sensors. There is growing indication that highlights the significance of the sampling and delivery location in wound care platforms that utilize the concept of "theranostics".

Microneedles (MN) have recently demonstrated significant promise in the development of multifunctional dressings for efficient drug delivery in CWOs therapy^[16]^. However, with microscale (typically 1 mm) needles, MN patches can readily penetrate skin and reach dermis level (where 40% of Interstitial fluid-ISF exits and WX is produced) without reaching nerves and blood vessels,^[17]^ allowing for minimally invasive on-site detections and therapies. ISF found in the cavities between cells in body tissues (the interstitium) is the source of WX. Interstitial fluid is formed from blood in capillaries and contains 83% of the same components as plasma^[18]^. The ISF transports cellular nutrients, signaling molecules, and metabolic detritus. It forms the basis of WX when it leaks into a wound cavity. The inflammatory process triggered by the formation of a wound releases a variety of substances (mediators and enzymes) that, among other effects, promote the formation of WX by increasing interstitial fluid production^[19]^. Finding a potential correlation between wound ISF and wound bed exudate could be a game-changing advance in the management of CWOs. The aforementioned core idea has frequently been overlooked in the field of wound care.

Biosensor technology based on MNs can be divided into two categories: on-needle sensing and off-needle sensing^[20]^. The electrodes for electrochemical measurement in the reported on-needle biosensors are solid and coated conductive MNs. The interaction between analytes and electrodes directly generates a measurable current response. Typically, these electrodes are not biocompatible and require expensive cleanroom fabrication^[21]^. Off-needle biosensors extract ISF for analysis and post-treatment. In these applications, hollow MNs made of glass, silicon, or metal are utilized to extricate ISF through capillary force^[22]^ which can be a drawback because their extraction volumes are limited. Hydrogel-forming MNs (HFMNs) are an alternative because they extract ISF into their swellable porous matrix, are capable of wound healing, and can maintain the wound environment’s hydration level.^[23][24]^ Despite these unique advantages, HFMNs have not yet been utilized for in-vivo and real-time sensing and therapeutics as for conductivity and stability problems. Due to the hydrophilic character of the expandable polymer, it is challenging to make the hydrogel electrically conductive.

It is critical to develop conductive hydrogels with superior electrical properties and aqueous stability when designing HFMNs for adverse chemical microenvironments (such as CWOs). Hydrogels that incorporate conducting 2D nanofillers and polymers (e.g., polystyrene sulfide, poly(3,4-ethylenedioxythiophene), rGO, MXene, and MoS_2_) exhibit considerable potential.^[25,26]^ On the contrary, the MXenes (Ti_3_C_2_T_x_), which are new two-dimensional transition metal carbides, exhibit enhanced electronegative functional groups (-OH, -F, -O), significant metallic conductivity, and a greater specific surface area. 3D polymer hydrogel networks could be synthesized with enhanced mechanical stability through the incorporation of electronegative groups of Ti_3_C_2_T_x_.^[27,28]^ In addition, MXene plays an important role in the healing of CWOs due to the antimicrobial activity of its surface functional groups^[29]^. Conductive hydrogel-based MNs composed of PEDOT: PSS have garnered the most attention due to their unique electrical and ionic dual conductivity and outstanding biocompatibility. However, when the PEDOT: PSS is mixed or embedded with the hydrogel, the electrical conductivity of the hydrogel decreases sharply (in the range of semiconductors).^[30]^ Moreover, PEDOT: PSS is unfavorable for long-term operation in contact with biological tissues due to its relatively high Young’s modulus (1 to 2 GPa), low stretchability (2% strain) due to the brittle PEDOT-rich domain, and water instability due to the hydrophilic PSS-rich domain.^[31]^ To convert PEDOT: PSS into water-stable soft hydrogels, phase separation techniques that redistribute the networks between the hydrophobic and conductive PEDOT-rich domain and the soft and hydrophilic PSS-rich domain have been developed.^[32,33]^ In addition, patterning and coating them with a high spatial resolution is a significant obstacle for applications involving microneedles. Recently, laser-induced phase separation of PEDOT: PSS introduced a potential method for devising a novel biocompatible and ultrafast digital patterning process for the fabrication of water-stable PEDOT: PSS hydrogels-based electrodes.

Here, we introduce a high-performance transdermal dressing system for the first time that uses a highly conductive, biocompatible, and minimally invasive replaceable microneedle array to monitor the physiological conditions of the transdermal wound ISF and performs combined therapy because of the hydrogel’s antibacterial and antimicrobial properties. Compared to previously reported wearable sensors that rely on wound bed exudate, theranostics at transdermal ISF can provide more comprehensive and customized information for effective chronic wound management. The HFMNs is made of polyvinyl alcohol, a biocompatible and swellable polymer, and chitosan, a crosslinking agent. Conductive nanofillers, such as MXene (Ti_3_C_2_T_x_) nanosheets, are added to the mixture to aid in the development of 3D polymer hydrogel networks by hydrogen bonding. The antimicrobial activity of the surface functional groups on MXene contributes significantly to the improvement of CWO healing and the enhancement of mechanical stability in hydrogels. Additionally, through the laser-induced phase separation of PEDOT:PSS/GO, we develop an innovative digital patterning method that is both biocompatible and capable of producing an highly conductive array of water-stable HFMNs. The LSPS significantly improved electrical conductivity and aqueous stability by virtue of the connected and enlarged PEDOT-rich regions. This theranostic approach establishes a correlation between wound bed exudate and wound-affected ISF, thereby facilitating the commercialization of the implantable dressing systems.

## 2. Results and Discussion

### 2.1 Design and Fabrication Strategies of the HFMNs-Based Dressing System

#### 2.1.1. Fabrication of Microneedle Array

Here, cleanroom-free stereolithography 3D printing was used to produce solid microneedles (Figure 1a, i). These microneedles were produced up of 49×43 conically-shaped needles (800 μm in height) arranged atop a 60×40 mm film (5mm in thickness), and they served as the master mold for the creation of polydimethylsiloxane (PDMS) negative molds (Figure 1a, ii and iii), which were then used for hydrogel casting on them (Figure 1b, i). Choosing to produce microneedle array using 3D printing can offer a scalable, reliably repeatable, and substantially less expensive production method.^[34]^ After two cycles of freezing and thawing, the HFMNs were extracted from PDMS, dried under vacuum, and used for further modifications. The hydrogel was made of polyvinyl alcohol (PVA), chitosan (Ch), and MXene (Ti_3_C_2_T_x_), in which the biocompatible PVA can form a 3D network with Ch via cross-linking by repeated freezing and thawing (Figure 1b, i), and Ch can improve the porosity and water absorption capacity of hydrogel.^[35]^ Due to the phase transition property and 3D network structure of the PVA/Ch hydrogel, the microneedle can easily penetrate the skin and extract an adequate amount of ISF. In addition to a larger specific surface area and metallic conductivity, MXene possesses an enhanced electronegative functional group (-OH, -F, -O).^[27]^ The introduction of electronegative groups onto Ti_3_C_2_T_x_ triggers the formation of three-dimensional polymer hydrogel networks via hydrogen (H) bonds (see Figure 1b, i inset). This mechanism improves the hydrogel’s mechanical and electrical properties. Dropcasting was utilized to deposit a 360° electroactive layer composed of PEDOT: PSS and graphene oxide (GO) onto both surfaces of the 3D-printed microneedles (Figure 1b, ii). This chemical method promotes the rapid mass production of conducting polymers onto the surface of a hydrogel in contrast to conventional electropolymerization techniques, which involve the formation of the conducting polymer onto a conducting substrate. After several washings and a thorough drying process, the composite-coated HFMNs were ready for the next phase.

**Figure 1.**
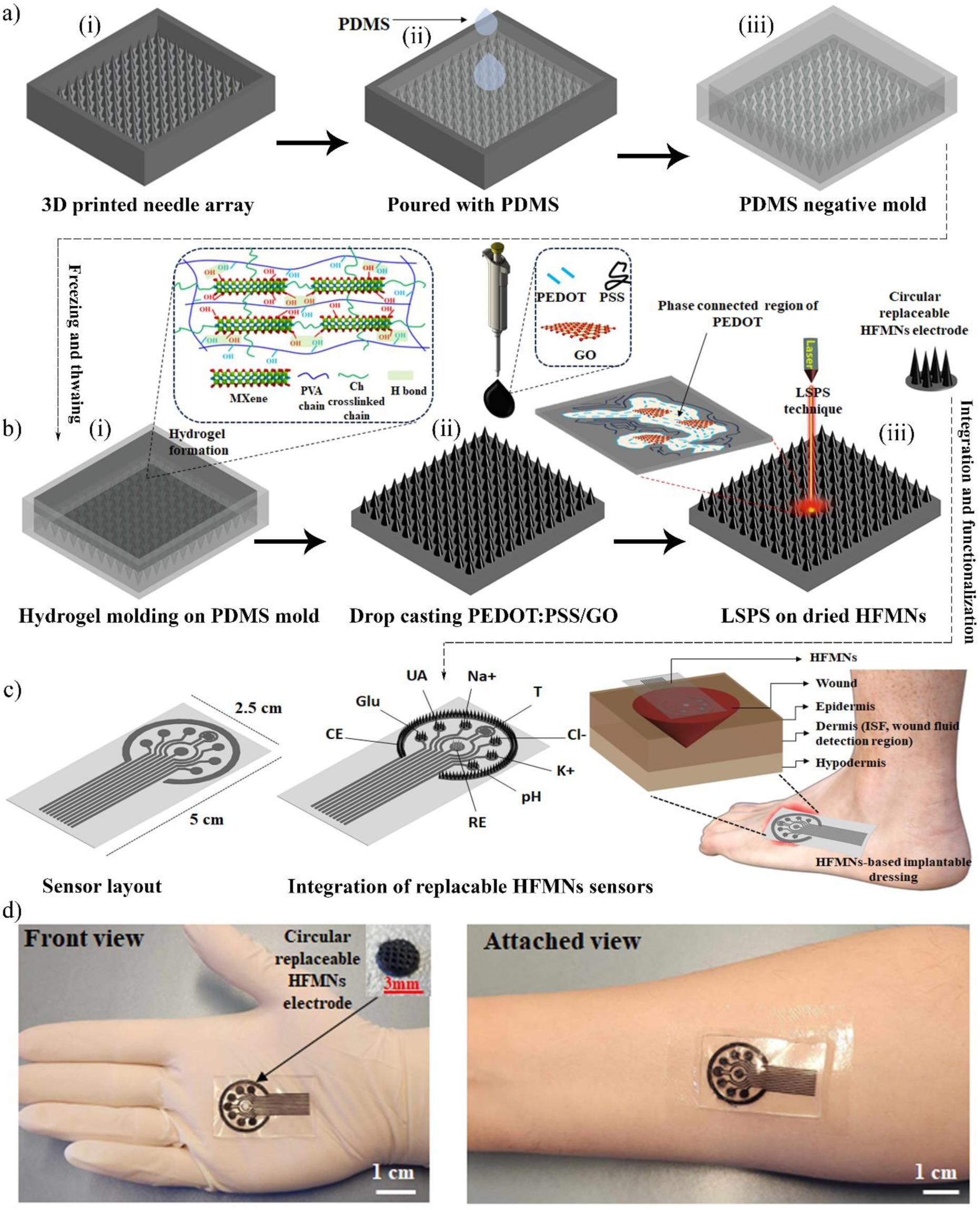
The fabrication concept of HFMNs array-based implantable dressing system for chronic wound diagnostics and therapy. a)(i) Poured the hydrogel on 3D printed negative mold. The hydrogel was made of polyvinyl alcohol (PVA), chitosan (Ch), and MXene (Ti_3_C_2_T_x_), in which the biocompatible PVA can form a 3D network with Ch via cross-linking by repeated freezing and thawing (inset), (ii) coating of PEDOT: PSS/GO, (iii) subsequent LSPS process and phase separation (inset). b) Fabrication of sensor layout on flexible substrate (CE-counter electrode and RE-reference electrode), integration and functionalization of the replaceable sensors onto the sensor pad, and placement of the HFMNs-based implantable system for CWs monitoring and therapy. c) Photographs of the fingertip-sized stretchable and flexible dressing system (front and attached view).

#### 2.1.2. Laser-Scribed Phase Separation (LSPS)

The conductivity of the composite hydrogel and its water stability are restricted by the loose extrinsic coating of conductive layer (PEDOT: PSS/GO), which poses challenges for the on-site sensing application and stability of HFMNs. PEDOT: PSS exhibits a phase dispersion in which the insulating PSS is located in the shell and the conductive PEDOT is located in the core. The PSS shell around the PEDOT core not only obstructs charge flow but also plays a role in PEDOT dispersion.^[36]^ A key approach for converting PEDOT: PSS into hydrogels that are water-stable is to restructure the phase organization of the material.^[37,38]^ Phase separation is a novel method for producing conductive PEDOT: PSS hydrogels by careful alteration of the orientation and crystallization of the PEDOT and PSS-rich regions, respectively. Precisely, outstanding electrical conductivity and aqueous stability can be achieved through a robust interconnection between the hydrophobic and conductive regions rich in PEDOT. Therefore, we employed the novel laser patterning process to fabricate highly conductive hydrogels (384 S/m) via the LSPS without the need for a separate postchemical cleaning step on PEDOT: PSS/GO coated HFMNs (Figure 1b, iii). In order to optimize laser absorption, PEDOT: PSS was sonicated with GO solution with varying concentrations as the solvent. This process facilitated the separation and recrystallization of PEDOT and PSS via a fast temperature increase and an enhanced electric field. We proposed that any laser is capable of simultaneously supplying photothermal energy and an electric field to PEDOT:PSS/GO, in light of the improved properties of PEDOT:PSS-based hydrogels ascribed to phase separation via green-laser scanning,^[38]^ dry annealing,^[39]^ and strong electric field^[40]^ published in prior research. The robust network between the PEDOT-rich regions enables a stable mechanical connection in water, while the expanded PEDOT region enhances the pathway for efficient charge flow.

#### 2.1.3. Fabrication of the Dressing System

After the LSPS, the desired circular-shape electrodes can be cut and utilized for replaceable sensors on a flexible, custom-designed polar sensor array layout (Figure 1c, i and ii) for CWs monitoring and treatment at the dermis level after functionalization. A transdermal dressing comprising a functionalized multiplexed microneedle array is illustrated in Figure 1c, iii. This dressing is utilized to apply on-site wound theranostics to open wounds in patients with CWs. The complete assembly was attached to a wearable platform using transparent, gas-permeable medical tape. The dressing substrates’ transparency makes it easy to view and assess the color and surface area of the wound. Figure 1d depicts a prototype of transparent transdermal dressing systems for CSO monitoring and treatment.

### 2.2. Morphological and Physical Characterizations

After the LSPS technique, HFMNs exhibited a conical structure with a length of approximately 800 μm, a pointed tip measuring around 10 μm, and an inter-needle spacing of 700 μm, as observed by scanning electron microscopy (SEM) (Figure 2a). This indicates that the coated laser-scribed microneedles possess uniform microstructural features after selective phase-separation. The phase separation characteristics of the laser-scribed hydrogel were examined in relation to the GO solution fraction in PEDOT: PSS. 2 mg/mL GO solutions (1–10 volume%) in water was sonicated with a composite PEDOT: PSS solution and dried. As we increased the GO volume %, the optimal laser power decreased, indicating that a greater GO volume fraction improved laser absorption and increased thermal and electric field effects. The relative concentration of PEDOT: PSS components, however, was inadequate to create a stable conducting network between the PEDOT-rich regions when an excessive quantity of GO was included. The robust expanded network connecting the PEDOT-rich region (Figure 2b) may facilitate a more efficient pathway for fast charge transfer. Atomic force microscopy (AFM) phase image analysis was used to verify the laser’s selective phase separation. Prior to laser scribing, the PSS-rich region (dark color) obscured the PEDOT-rich domain, which was previously small and bright in color (Figure 2c). After the LSPS, the PEDOT-rich domain significantly increased and interconnected (Figure 2d). When the laser was working properly, increasing the laser power (LP) made PEDOT: PSS more electrically conductive and stable in water. But when the laser power was too high, PEDOT: PSS turned into carbon and its conductivity started to drop (Figure 2e). High speed laser scribing (300 s) may obtain the maximum electrical conductivity (sheet resistance of 13 Ω/Sq) for LP 12% when combined with the impact of GO.

**Figure 2.**
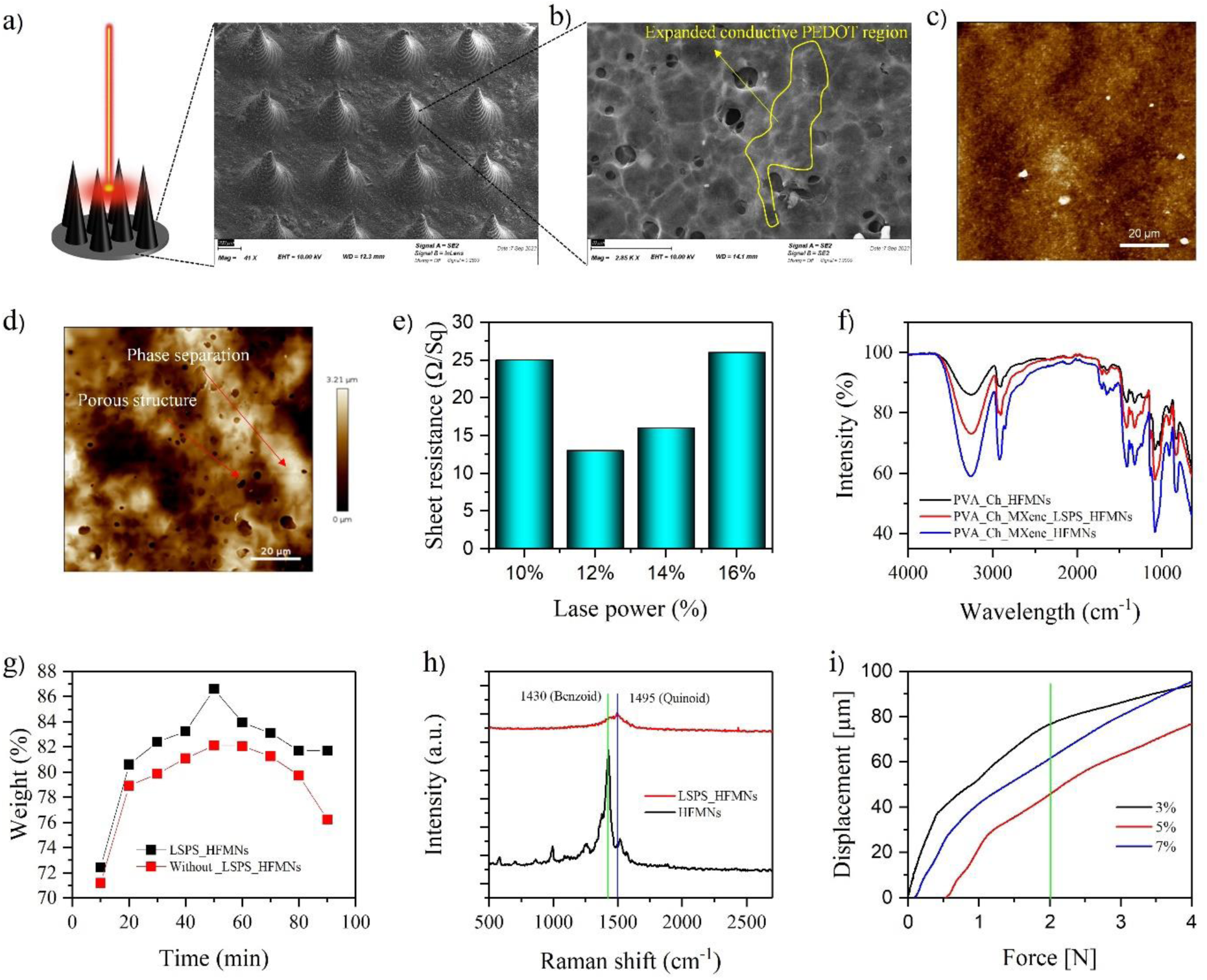
Physical characterizations of the HMNs a) SEM image of the LSPS HFMNs. The array exhibited a conical structure with a length of ∼800 μm, a sharp tip of ∼10 μm, and inter-microneedle spacing of 700 μm after the LSPS technique. b) Expanded porous PEDOT rich region due to LSPS. c) and d) AFM phase-separation image of HFMNs. The PEDOT-rich domain was greatly expanded (bright color) and connected after the LSPS. e) Laser power optimization for LSPS. For LP 12%, combined with the effect of GO, fast laser scanning (300 s) can achieve the highest electrical conductivity (sheet resistance of 13 Ω/Sq). f) FTIR spectra analysis. PVA/Ch, PVA/Ch/MXene, and LSPS-based PVA/Ch/MXene HFMNs showed peaks at 3417, 2927, 1616, 1413, and 1029 cm^-1^, which correspond to the O-H, C-H, COO-, (C═O), and C-O-C vibration. g) Swelling ratio. The results showed that the LSPS had a high effect on the water content of the gel and increased its swelling rate and stability due to its porous morphology. h) Raman analysis. i) Mechanical chraracterizaion.

The chemical bonding of the hydrogel is confirmed by Fourier transform infrared spectroscopy (FTIR), as illustrated in Figure 2f. The peaks at 3417, 2927, 1616, 1413, and 1029 cm^-1^ in the FTIR spectra of the PVA/Ch, PVA/Ch/MXene, and LSPS-based PVA/Ch/MXene HFMNs correspond to the stretching vibrations of O-H, C-H, COO-, C=O, and C-O-C, respectively.^[29,41]^ It provides evidence that supramolecular H bonding networks are formed between PVA, Ch, and Ti_3_C_2_T_x_. Comparatively, the maximal intensity of HFMNs appears to have decreased marginally following LSPS as a result of an increase of hydrophobic PEDOT-rich regions.

In order to evaluate the impact of LSPS on the swelling capacity of HFMNs, their mass changes over time were observed in phosphate buffered saline (PBS). The swelling ability of HFMNs is determined by their porous morphology. The findings depicted in Figure 2g indicate that the LSPS significantly impacted the gel’s water content and enhanced its swelling rate as a consequence of its porous structure. Notably, the LSPS method enhances the aqueous stability of the hydrogel through the conversion of hydrophobic PEDOT-rich regions.

Additionally, Raman spectroscopy was employed to examine the molecular configuration of the PEDOT-rich region (Figure 2h). The PEDOT crystallite possesses a benzoid structure (1430 cm^−1^) characterized by its relatively low electrical conductivity, as well as a quinoid structure (1495 cm^−1^) that is associated with its high electrical conductivity.^[42,43]^ This conformation shift in the PEDOT morphology from a helical structure to linear structure—also known as the secondary doping effect— was suggested by the observed decrease in benzoid and rise in quinoid structure.^[43]^

Subsequently, mechanical integrity testing was conducted in order to ascertain whether various compositions of HMN possess sufficient compressive strength to penetrate the epidermis effectively. A minimum force of 0.045 N/Needle is necessary in order to achieve epidermal piercing.^[44]^ When tested under mechanical compression, none of the microneedles exhibited any indications of plastic distortion or failure, as evidenced by the force-displacement and stress-strain relationship that was nearly linear (Figures 2h). An increase in MXene concentration (from 3% to 7%) resulted in a displacement of 76 μm for 3% HFMNs at 2 N. However, for 5% MXene, the recoded displacement decreased to 45 μm, indicating an enhancement in mechanical rigidity. Nonetheless, an excess of MXene 7% induced an increase in displacement of 61 m, which might be attributed to agglomerated 3D hydrogel formation. When compared to other hydrogel microneedles, our results indicated exceptional mechanical _stability._^[30,44,45]^

### 2.3. Characterization of the HFMNs sensor array for multiplexed CWO-biomarkers analysis

We employed circular replaceable array sensors as an efficient method to extend the operational duration of the implantable systems, accounting for inevitable deactivation of enzymes and mechanical friction. A polar-type sensor array of seven electrodes was engineered utilizing laser-scribed graphene-based flexible electrodes (2.5×5 cm) with their connections passivated by a waterproof layer of plasticizer. All active circular laser-cutter HFMNs-based sensors [glucose (Glu), uric acid (UA), pH, Na+, Cl-, K+, and temperature (T)] along with the counter electrode (CE) and reference electrode (RE) are positioned in separate and replaceable modules. Hence, a replacement array may be quickly installed in the sensor platform in case that an electrode is used and not working. Additionally, in a manufacturing setting, it will be simpler to change the production line from one analyte to another. Following the brush-painting of a thin layer of conductive material as the liquid adhesive onto the electrode pad, an array of functionalized HFMNs is applied for further characterizations. Following the brush-painting of a thin layer of conductive material as the liquid adhesive onto the electrode pad, an array of functionalized HFMNs is applied. Artificial ISF was utilized to assess the sensor’s performance. We implemented this innovative approach to produce implantable dressing systems composed of HFMNs for the first time. The concentrations of critical indicators in WX and ISF might be considered as (glucose: ISF (0-7) mM and WX (0-28) mM, UA: ISF (200-600) µM and WX (0-760) µM, Na^+^ and Cl^-^: ISF: (0.75-200) mM and WX (5-160) mM, K+: ISF (1-128) mM and WX (3.2−5.7) mM, pH: ISF (5-9) and WX (4-9), and temp: (25-40)°C.^[46,47]^ This results in a significant challenge in achieving linear sensor response in physiological concentration ranges, as the concentration range of the corresponding markers in ISF is greater than that of wound exudate. In order to mitigate these concerns and obtain precise monitoring of the metabolism, we utilized a novel functionalization process and optimized diffusion liming membrane. We engineered our enzymatic Glu and UA sensors using platinum nanoparticles/EDC-NHS/Glucose oxidase or Uricase (PtNPs/EDC-NHS/GOx or UOx) with extra polyurethane (PU) permeable membrane coatings. The triple coating of PU-based enzymatic sensors demonstrated the best linearity and repeatability in a complicated fluid matrix. The enzymatic glu and UA sensors showed excellent selectivity, and their amperometric responses were proportionate to the relevant concentrations of the corresponding markers in ISF, with linearities of 0.45–31.45 mM and 0.15-2.75 mM and sensitivities of 12 and 80 µAmM^−1^cm^-2^, respectively. (Figure 3, a to d). For Na^+^, Cl^-^, and K^+^ sensors, the response mechanism relies on how changes in ion levels alter the associated ion-selective membrane potentials. Figures 3e-i show the open circuit potential responses of the Na^+^ (1-366 mM), Cl^-^ (1-366 mM), and K^+^ (1-156 mM) in ISF solution. Excellent linearity was observed in all three ion-selective electrodes, demonstrating average sensitivities of 48 mV for Na^+^, 41 mV for Cl^-^, and 32 mV for K^+^ per decade of concentration, which are also close to the slope of the Nernst equation. All of the ion sensors demonstrated high selectivity against various interferents (Figures 3g and 3i). The sensor array includes an integrated PtNPs-based resistive T sensor. In a typical physiological range of 25° to 40°C, it has a sensitivity of approximately 0.35% °C^-1^ (Figure 3j). Figures 3k and 3l illustrate that the pH sensor, which employs an electrodeposited polyaniline layer as the pH-sensitive membrane, demonstrated a good selectivity and sensitivity of 45.7 mV pH^-1^.

**Figure 3.**
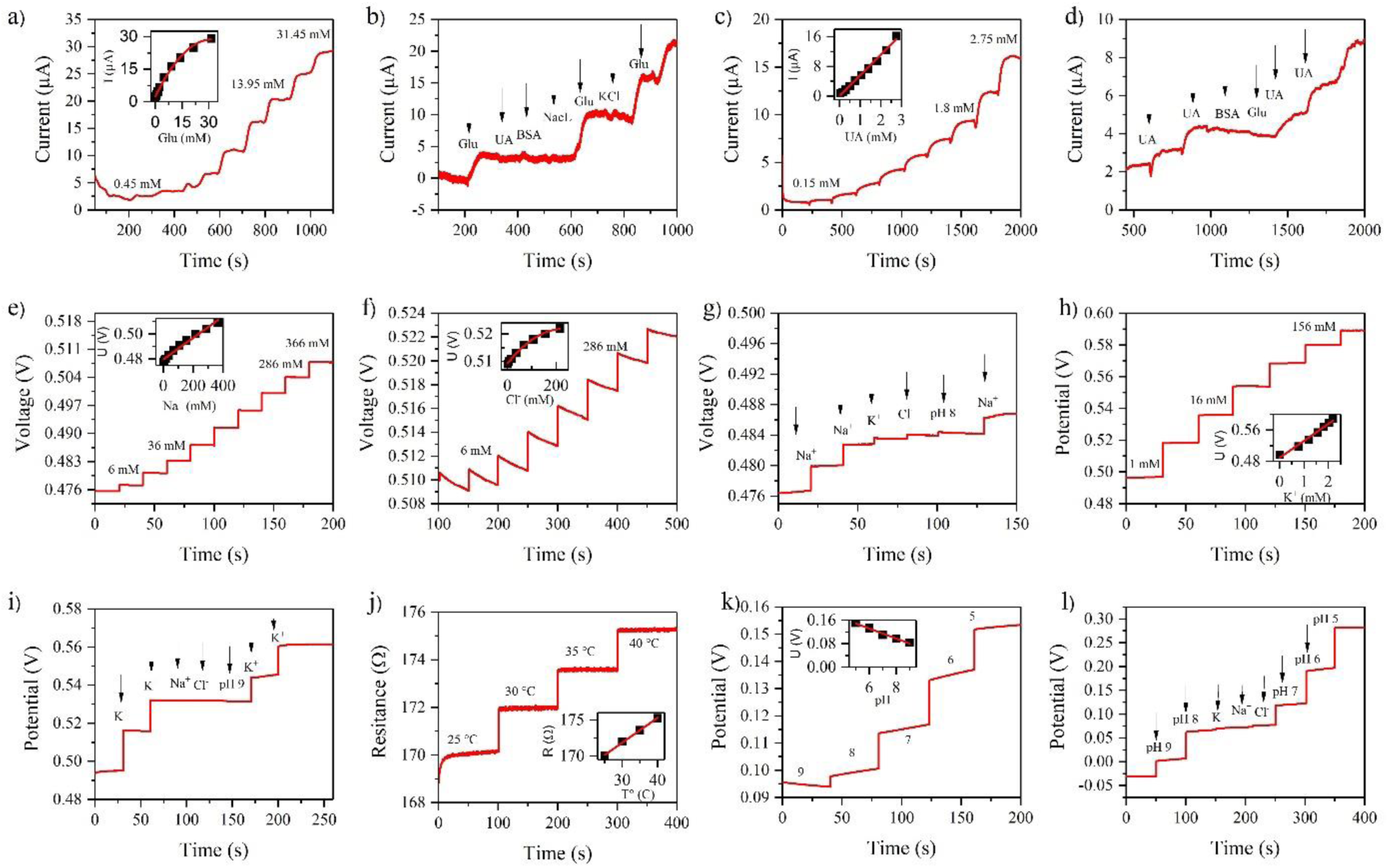
Characterization of a sensor array for multiplexed biomarkers analysis in ISF. (a-d) glu and UA selectivity and amperometric responses. Calibration maps with a linear fit are insets in b) and d). e-i) Potentiometric response to Na^+^,Cl^-^ and K^+^. Calibration plots with a linear fit are insets in e) and g). g) and i) Selectivity tests of the ion sensors. The physiologically relevant temperature range induces a resistive response in a Pt-based T sensor. Insets, the linearly fitted calibration plot. k) Open circuti potential response of a pH sensor based on polyaniline. With a linear fit, insets depict the calibration plot. l) Selectivity test of the pH sensor.

### 2.4. Correlation Between Wound ISF and Exudate

As previously determined by the HFMNs-based implantable system, the biomarkers were subsequently evaluated independently in patients’ artificial wound exudate utilizing conventional methods in order to validate and establish the correlation with the developed implantable system. Figure 4a-g depicts the outcomes of monitoring each indicator in three distinct states (pre-infection, post-infection, and healing state) in ISF and exudate through the use of conditions that closely resemble the actual states. Considerable increases in UA, T, and pH were detected in comparison to pre-infection levels. ^[15]^ There is a potential correlation between the rise in temperature and inflammation.^[48]^ An activation of xanthine oxidase, an element of the innate immune system that is involved in the production of UA through purine metabolism in response to inflammatory cytokines in wounds, may account for the increased concentrations of UA following infection.^[49]^ Because of their relationship to acidity, pH and UA have both been shown to rise during bacterial infections.^[50]^ On the contrary, the glucose concentration in wound fluid that was infected decreased by more than 36% subsequent to the infection. This decline can be attributed to the heightened consumption of glucose by the bacteria.^[51]^ After treatment, the glu level considerably increased, while the T, pH, and UA returned to their pre-infection values, signifying the effective removal of the bacteria. Similarly, during the initial phases of infection, significantly high levels of Na^+^ and K^+^ were observed. Next, the levels of Na^+^, Cl^-^, and K^+^ in the exudate gradually dropped and then showed an increasing trend once more as the wound healed. For a normal wound, this could be explained by the need for excess ions to be involved in the healing process, whereas for an infected wound, it could be linked to bacterial growth.^[8]^ The ISF measurements displayed comparable results to those of the exudate, indicating that the HFMNs are capable of providing objective quantitative data that fits within clinically significant ranges.

**Figure 4.**
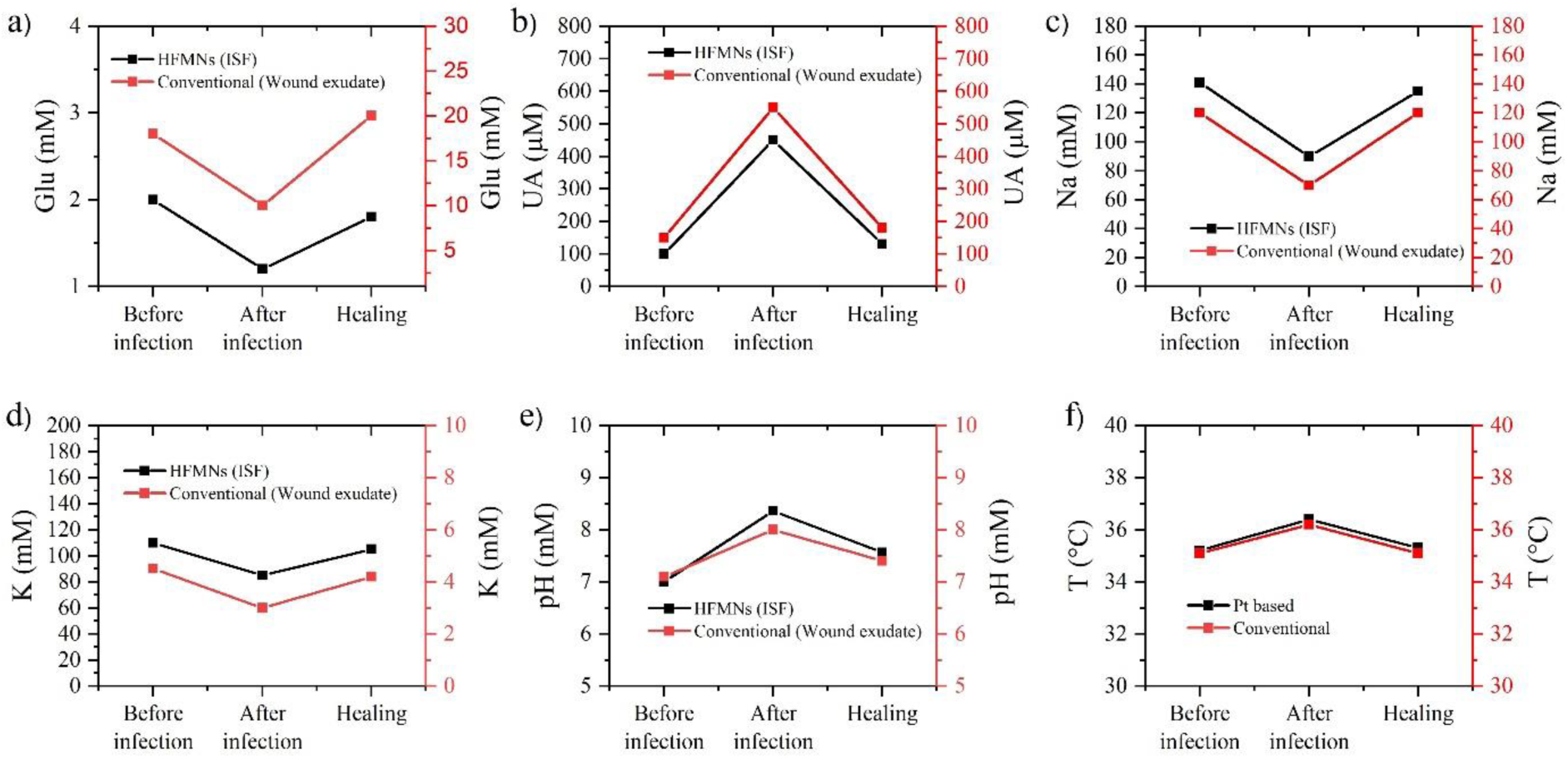
Correlation of the HFMNs system and conventional sensors for multiplexed wound biomarker monitoring in wound ISF and exudate, respectively, in three different ex vivo systems (before infection, after infection, and healing). a-f) Glu, UA, Na^+^, K^+^, pH, and T sensors.

### 2.5. In-vitro Antibacterial Activity and Therapeutic Capabilities

Strong antimicrobial properties are essential for the perfect wound dressing. Bacterial enumeration was employed to ascertain the antibacterial activity of the HFMNs against *E. coli* and *S. aureus*. These bacteria were incubated at 37 °C for 6 hours with PVA, PVA/Ch/MXene, and LSPS-based PVA/Ch/MXene HFMNs, as illustrated in Figure 5. As a result of the bacterial cell membrane-attacking antibacterial activity of MXene’s surface functional groups, the PVA/Ch/MXene group contained fewer bacteria than the PVA group, and its antibacterial effect was substantially enhanced via the LSPS process. It is worth mentioning that the application of the LSPS process resulted in a substantial reduction in bacterial count within the PVA/Ch/MXene hydrogel treatment group, with a mortality rate exceeding 95% (survival rate of *E. coli* 1.66 and *S. aureus* 10). The favorable antibacterial outcome can be primarily ascribed to the synergistic interaction between the exceptional photothermal effect of MXene on LSPS and its intrinsic antibacterial characteristics.

**Figure 5.**
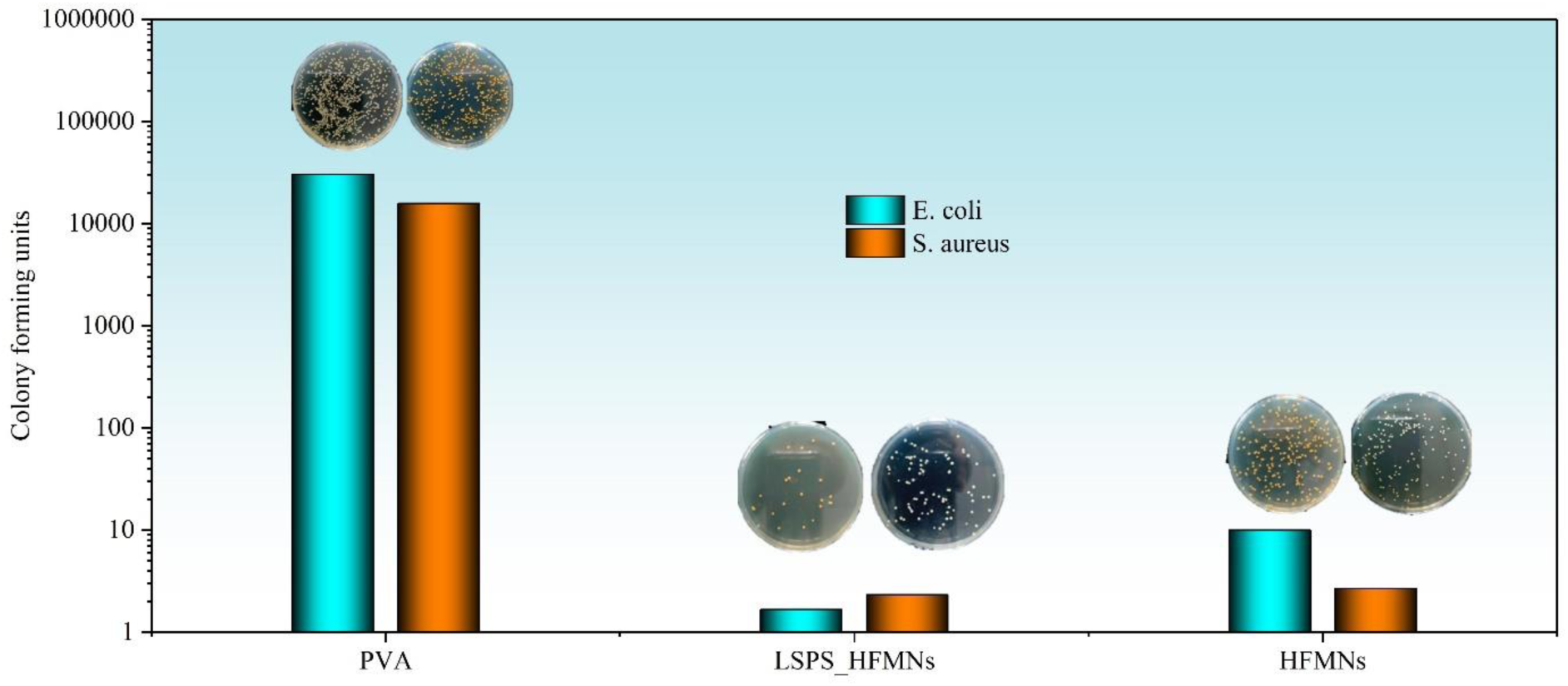
Antimicrobial studies. Expreiments and photograph of bacterial colonies of E. coli and S. aureus treated with hydrogels containing different compositions: PVA, HFMNs, and LSPS_HFMNs.

## Conclusion

We introduce the first transdermal dressing systems comprised of highly conductive HFMNs that enable antimicrobial healing and multiplexed monitoring of wound-related biomarkers from dermis-level ISF. The HFMNs exhibit remarkable stretchability, mechanical stability, and complete biocompatibility. Moreover, it possesses the ability to selectively monitor a panel of vital wound biomarkers in real-time, including glucose, uric acid, pH, Na^+^, Cl^-^, K^+^, and temperature. Furthermore, we present a methodology for the LSPS process-based patterning and production of a high-performance microneedle array. The produced hydrogels exhibited exceptional electrical characteristics and maintained their electrical and electrochemical capabilities in an aqueous environment for an extended period of time. The implanted dressing system serves as a flexible platform for assessing wound conditions and providing intelligent therapy. It is also readily reconfigurable to monitor a number of other metabolic and inflammatory indicators for a range of applications related to chronic wound care. Our outcomes and the correlation between wound ISF and exudate suggest that wound fluid analysis and treatment provided by implanted dressings may be a viable means of achieving ongoing and customized wound care. However, the lack of in-vivo applications, which we intend to incorporate into the university clinic shortly, is a limitation of the current study. We foresee the fully integrated, custom-engineered dressing system functioning as a platform for personalized monitoring and treatment of chronic wounds that is easier to implement, more effective, and completely controllable.

## Experimental Section

### Materials and Regents

MAX phase (Ti_3_AlC_2_) (≥40 μm) powder, potassium tetrachloroplatinate, cyclohexanone, tetrahydrofuran, bis(2-ethylhexyl)Sebacate (DOS), N-Hydroxy succinimide (NHS), N-(3-Dimethylaminopropyl)-ethylcarbodimide hydrochloride (EDC), polyurethane, graphene Oxide, poly(3,4-ethylenedioxythiophene)-poly(styrenesulfonate)(PEDOT:PSS), aniline [C_6_H_7_N], chitosan, D-glucose , sucrose, uric acid, poly vinyl chloride (PVC), poly vinyl alcohol (PVA), glucose oxidase from Aspergillus Niger, uricase from Candida Sp., sodium-(triflutomethyl)Phenylborate[Na-TFPB], bovine Serum albumin (BSA), selectophore grade sodium ionophore X, sodium tetrakis[3,5-bis(trifluoromethyl)phenyl] borate (Na-TFPB), valinomycin (potassium ionophore), sodium tetraphenylborate (NaTPB), dimethyl sulfoxide (DMSO), potassium ferricyanide (K_3_Fe(CN)_6_), HCl, H_2_SO_4_, glutaraldehyde solution (20−25%), poly(vinyl butyral), sodium chloride (NaCl), potassium chloride (KCl), calcium chloride (CaCl_2_), sodium bicarbonate (NaHCO_3_), sodium phosphate (NaH_2_PO_4_), magnesium sulfate (MgSO_4_), sodium gluconate (NaC_6_H_11_O_7_), lactate, buffer solution (pH = 4, 7, and 10), PBS (pH = 7.2), and Sylgard 184 elastomer kit (PDMS and curing agents) were bought form Sigma-Aldrich (Germany). All solutions were prepared using deionized water produced from a Millipore water purification system, unless otherwise noted.

### Fabrication of the Microneedle Array

AutoCAD was utilized in the development of the computer-aided design (CAD) file. The CAD was comprised of a circular base measuring 60×40 mm film (5mm in thickness), which featured 49×43 conically-shaped needles with an interneedle spacing of 700 μm and a base diameter of 300 μm and a height of 800 μm, respectively. Following its exportation as a standard tesselation language (.STL) file, the document was imported into the Preform 3D printing application developed by FormLabs, USA. Prior to generating supports with a reduced touchpoint size and configuring the high temperature resin to print at a high resolution, the component was initially inclined at an angle of 45°. After the printing process, the resulting prints underwent a 60-minute washing in IPA with FormWash, followed by a 150-minute treatment at 70 °C under 405 nm UV with FormCure. A scalpel was employed to remove the supports by means of a gentle break at the interface where the supports met the base of the microneedle array.

### Fabrication of HFMNs

HFMNs involves total four vital phases: i) preparing the homogenous hydrogel solution, ii) pouring it into a PDMS female mold, iii) cross-linking by freezing followed by thawing, furthermore, iv) drying and demolding the microneedle patch. In first phase, 15% (w/w) PVA %solution was prepared with addition of 3%, 5%, and 7 % MXene solution, heated at 90°C for 3 hours. For the synthesis of MXene, we will utilize the modified minimally intense layer delamination MILD synthesis method, as described in our prior publication.^[52]^ Separately, 1% (w/w) Ch solution was prepared in 0.1M acetic acid at room temperature (RT). Subsequently, PVA/CS solutions ratios (4:1) were mixed, and stirred overnight at RT. The PVA/CS solution was subsequently injected into a PDMS female mold applying a vacuum to fill the microneedle pores in the second phase. Furthermore, the hydrogel forming microneedles (HFMNs) was frozen at −20°C for 3 hours and thawed at room temperature, then dried in oven at 50°C for 6-8 hours. Afterwards, the microneedles patch was removed carefully from PDMS mold.

A GO solution (2 mg/mL GO) was sonicated with PEDOT: PSS aqueous solution at a concentration of 1–10 volume% in water. The final ink was applied to the HFMNs array after 1-hour of sonication at room temperature. The array was then allowed to dry for 24 hours at RT.

The laser system comprised a 10.6 µm CO_2_ that was employed to scribe selective phase separation via laser processing. For a final PEDOT: PSS containing 10% GO, the optimal conditions in this investigation were 12% laser power and 300 mm/s scanning speed. Following laser treatment, DI water was used to rinse the samples.

### Fabrication of the Transdermal Dressing System

The methodology for producing the electrode arrays is illustrated in Figure 1. Using AutoCAD, the distinctive polar base electrode patterns were designed. We subsequently patterned the 3D porous laser-induced graphene-enabled patterns onto commercial PI films using the CO2 laser scribing technique (optimized parameters: power: 14%, speed: 250 mm/s). A vacuum-degassed PDMS solution (base: curing agent = 10:1) was subsequently poured onto the scaffold containing the pattern. The pattern was carefully picked up by the PDMS film following a successful two-hour thermal curing process at 70 °C. Next, SEBS was employed for passivation. The patch was then transferred to a transparent and breathable medical-grade tape (Tegaderm) and applied to the subject’s wound region.

A PalmSens potentiostat was utilized to assess the linear range, sensitivity, and reproducibility of the multiplex sensors in solutions of target in artificial ISF and synthetic wound fluid. The synthetic ISF and wound fluid were prepared in accordance with the methodology previously described.^[15,53]^

### Fabrication of the Replaceable Sensors

In accordance with a methodology, we have previously established, Pt NPs were electrodeposited onto glu and UA sensing HFMNs.^[52]^ A Pt NPs precursor solution was utilized to immerse the circular HFMNs, which were subsequently electrodeposited at a scan rate of 50 mV s^−1^ for 25 segments using a CV ranging from -0.1 to 0.8 V. Utilizing the NHS/EDC mixture, the functional groups were activated. The array was drop-coated with a solution consisting of EDC and NHS (10 mM). Following this, the array was incubated at RT for a duration of 12 hours.

Following this, a solution containing 5.05 mg chitosan (acetic acid and DI) was combined with 10 mg GOx and UOx. Following 5 min stirring, 10 µL of the solution was dropped onto the HFMNs surface and dehydrated at 4 °C for a duration of 24 hours in a refrigeration unit. The working electrode was subsequently coated with PU (1%, 2%, and 5%) in the capacity of a porous membrane. Following a 20-minute drying period at RT, the sensor was stored at 4 °C until it was utilized. The HFMNs electrode was coated with Ag/AgCl by hand, which was then cured at 120 °C for five minutes in order to produce reference electrodes for the glu, UA, and pH sensors. In their unaltered state, HFMNs were utilized as the counter electrode.

Membranes for ion sensors were produced in accordance with an earlier report. The components of the Na+ selective membrane mixture were DOS (65.45%), PVC (33%), Na-TFPB (0.55%), and Na ionophore X (1%). Tetrahydrofuran was used to dissolve 2 ml of membrane cocktail (300 mg). The K+-selective membrane contained valinomycin (2% w/w), NaTPB (0.5%), PVC (32.7% w/w), and DOS (64.7% w/w). A solution of 300 mg of the membrane mixture and 700 μL of cyclohexanone was prepared. Ag NPs were deposited for the Cl^-^ ion membrane, and Fecl_3_ was subsequently employed to chlorinate it. The HFMNs were subsequently treated with ion-selective membranes developed by drop-casting 8 μL of the Na^+^ and K^+^ selective membrane mixture.

The HFMNs electrode was immersed in PANI precursor (0.1 m aniline aqueous solution with 1 m HCl) in order to electropolymerize the pH sensor. Then, 50 segments of CV were conducted at a scan rate of 100 mV s^−1^. The peak currents following electropolymerization increased rapidly with the number of deposited segments, indicating the successful coating of PANI membrane . Pt NPs were deposited and an interdigitated heater array was used to create the T sensor.

### Electrical Measurement

The sheet resistance of hydrogels was measured using a four-point probe. Using the thickness data acquired earlier, the electrical conductivity was calculated with the following equation

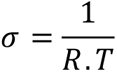

Where, σ, R, and T are conductivity, sheet resistance, and thickness.

### Mechanical Properties and Penetration of HFMNs

The dynamic mechanical analyzer (DMA) was employed to assess the mechanical properties of the conductive HFMNs. An upper titanium plate was lowered until contact was made with the microneedle points, as indicated by a force reading on the apparatus screen, while the microneedles were affixed to the lower titanium plate. The displacement produced when a compressive force ranging from 0 to 8 N was issued from the bottom plate at a rate of 0.20 N min^−1^ was measured. To observe the impact of compression on the microneedles, images were captured of the samples subsequent to the testing process.

### In-vitro Antimicrobial Studies

Antimicrobial testing was performed in Institute for Infection Medicine, Kiel, Germany according to previous report.^[54]^ In brief, 10 μL of an overnight culture of Escherichia (E.) coli and Staphylococcus (S.) aureus were incubated with 5 mL LB medium at 37°C for three hours. Subsequently the bacteria number was estimated by measuring the optical density at 600 nm. 10 µl of a suspension of 10^5^ bacteria/ml in 0.85% NaCl and 1% LB-medium were placed on the surface of the PVA membranes. The membranes were incubated in Petri dishes at 37 °C. After 6 hours, the membranes were transferred into 1 mL 1% LB medium with 0.85% NaCl and vortexed for 30 seconds. Serial dilutions of the supernatant were plated on LB agar und resulting bacterial colonies were counted to determine the number of surviving bacteria.

## Acknowledgements

Z.A. thanks the Schleswig Holstein State Fonds and Aventis Foundation (Grant number: 80304368) for the financial support.

